# Age-specific prevalence of anti-Pgp3 antibodies and severe conjunctival scarring in the Solomon Islands

**DOI:** 10.1101/141135

**Authors:** Robert Butcher, Oliver Sokana, Kelvin Jack, Diana L Martin, Matthew J Burton, Anthony W Solomon, David CW Mabey, Chrissy h. Roberts

## Abstract

**Background:** Trachomatous trichiasis (TT) and ocular *Chlamydia trachomatis (Ct)* infection in the Solomon Islands are scarce, whereas trachomatous inflammation–follicular (TF) is prevalent.

**Methods:** We enrolled 1511 people aged ≥1 year from randomly selected households in 13 villages in which >10% of the population had TF prior to a single round of azithromycin MDA undertaken six months previously. Blood was collected from people of all ages to be screened for anti-Pgp3 antibodies. Photographs were collected from people of all ages for analysis of scarring severity.

**Results:** Conjunctival scars were visible in 13.1% of photographs. Mild (p<0.0001) but not severe (p=0.149) scars increased in prevalence with age. Anti-Pgp3 antibody seroprevalence was 18% in 1–9 year olds, increased sharply around the age of sexual debut, and reached 69% in those over 25 years. Anti-Pgp3 seropositivity did not increase significantly between the ages of 1–9 years, and was not associated with scarring in children (p=0.472) or TF in children (p=0.581).

**Conclusions:** Signs of trachoma are common in the Solomon Islands but occur frequently in individuals who have no serological evidence of prior ocular infection with *Ct.* WHO recommendations for directing MDA provision based on signs alone may not be suitable in this context.

## Introduction

Trachoma is responsible for approximately 1.9 million cases of visual impairment or blindness globally.^1^ International partners have committed to elimination of trachoma as a public health problem by the year 2020. Repeated infections with *Chlamydia trachomatis (Ct*) and the immunological response to them can cause a gradual accumulation of scar tissue in the tarsal conjunctivae^2^. Scarring typically begins to develop in late childhood^3^ and can reach a prevalence of 25% in 10-year-olds in hyperendemic populations^4^. Scarring progresses throughout a lifetime and, in some cases, can lead to entropion, trichiasis, abrasion of the cornea, corneal opacity and blindness^5^. Infection with *Ct* induces the polyclonal production of antibodies to *Ct* antigens^6–8^. Measuring population-level accrual of antibodies to *Ct* is under investigation as a tool to monitor transmission of both urogenital^9^ and ocular^10^ infections.

Global trachoma elimination strategies are guided by the signs trachomatous trichiasis (TT) and trachomatous inflammation—follicular (TF). The World Health Organization (WHO) recommends at least three years of mass drug administration (MDA) with azithromycin in districts with >10% TF prevalence in 1–9 year-olds^11^. As trachoma control reduces prevalence, the positive predictive value of TF as a *Ct* infection marker drops and phenotypically similar diseases with other aetiologies will be unmasked.^12^ We have previously reported data from a 2013 population-based prevalence survey covering the two provinces of Temotu and Rennell & Bellona of the Solomon Islands, which showed the prevalence of TF in those aged 1–9 years was moderately high (26.1% of those examined), but TT (0.1% of those examined), trachomatous inflammation—intense (TI; 0.2% of 1–9 years-olds examined) and ocular infection with *Ct* (1.3% of 1–9 year-olds examined) were rare^13^. In accordance with WHO guidelines, MDA took place throughout the Solomon Islands in 2014. The program administered approximately 24,000 doses of azithromycin and achieved coverage of approximately 80% in Rennell & Bellona, and 85% in Temotu. After the baseline trachoma mapping in the Solomon Islands, the National Program also used billboard posters and regular radio spots to promote facial cleanliness and raise awareness of trachoma elimination.

The discordance between TF prevalence and *Ct* infection, TI and TT prevalence led us to question whether the observed TF was of chlamydial aetiology. Antibodies to *Ct* and trachomatous scarring (TS) are less susceptible to temporal variation than transient markers such as TF or infection and may be more informative of an individual’s cumulative trachoma history. We set out to determine whether the high TF prevalence in Rennell & Bellona and Temotu provinces was concurrent with significant burden of TS and whether TF occurred exclusively in those who had previously been infected with *Ct.* To address these questions, villages that previously had high proportions of children with TF were revisited 6 months after MDA and specimens were collected for assessment of anti-Pgp3 antibodies and scars.

## Methods

### Ethics

Study approval was from the London School of Hygiene & Tropical Medicine (LSHTM; 8402) and Solomon Islands National Health Research Ethics Committee (HRC15/03). Subjects aged 18+ years gave written informed consent to participate. A parent/guardian provided consent for those aged under 18 years.

### Study design

This study took place in June–July 2015, 6 months after a single round of azithromycin MDA had been delivered by the Solomon Islands National Trachoma Elimination Program. To enable comparison to pre-MDA data, only villages in Temotu and Rennell & Bellona provinces where baseline mapping had been conducted were eligible for inclusion. Thirteen villages where over 10% of the community (all ages) had signs of TF^13^ were selected. Due to their small respective populations (Temotu: 21,362; Rennell & Bellona: 3041), the two provinces were combined into one evaluation unit during baseline mapping. The number of villages in each province surveyed as part of this study were selected to reflect the relative population proportion (Temotu: 11 villages; Rennell & Bellona: 2 villages).

This survey was powered to estimate the prevalence of anti*-Ct* antibody seropositivity in children aged 1–9 years. Based on the low prevalence of ocular *Ct* infection prior to MDA (1.3%), we expected the seroprevalence to be approximately 10%, in line with other communities with low *Ct* prevalence^14^. To estimate seroprevalence with ±5% precision at the 95% confidence level, 367 children were required^15^. In our 2013 survey, we examined a mean of 1.1 children per household and therefore needed 25 households in each of 13 clusters to reach our sample size. All residents aged 1 year or above living in households drawn at random from a list of all households in a study cluster were eligible to participate.

### Trachoma grading

Grading using the WHO simplified system^16^ for TF, TI and TT was performed in the field by one of two Global Trachoma Mapping Project (GTMP)-certified graders, wearing 2.5× binocular magnifying loupes^17^. High-resolution digital photographs of the right tarsal conjunctivae were graded for TS using the modified WHO trachoma grading system (Table 1)^18^. Photographs were graded by two photo-graders who had previously achieved kappa scores for inter-observer agreement of >0.8 for F, P and C (follicles, papillae and cicatricae) grades, compared to a highly experienced trachoma grader. Photograph grading was undertaken masked to field grading, laboratory results, and the other photograph grader’s assessment. Discrepant grades were arbitrated by a third highly experienced grader.

**Table 1.**
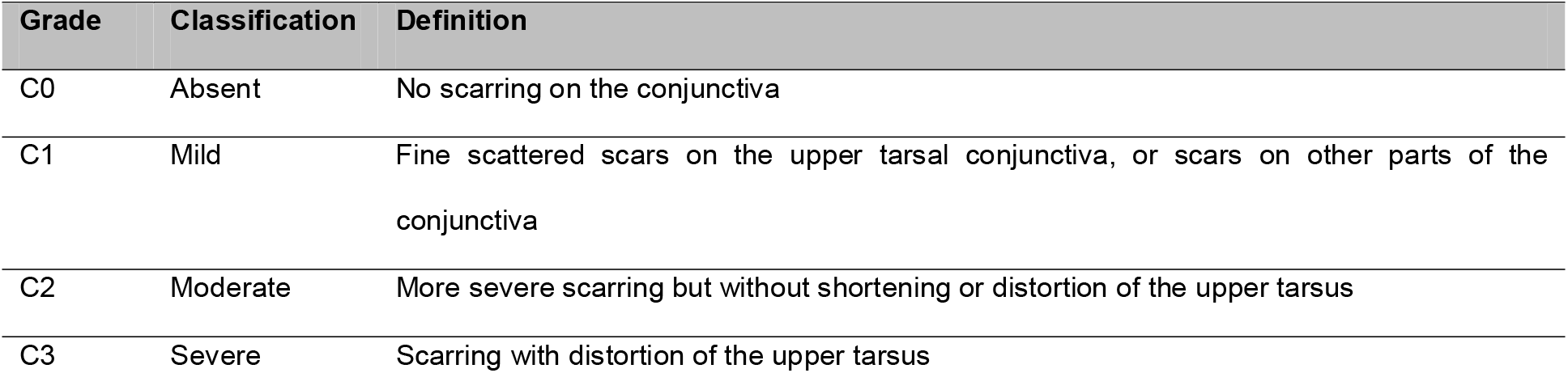
Grading system for conjunctival scarring, reproduced from Dawson and colleagues ^18^

### Specimens

Dried blood spots were collected for assessment of anti-Pgp3 antibody titre. Participants’ fingers were pricked with sterile lancets (BD Life Sciences, Oxford, UK) and 10 μL of blood was collected onto filter paper (CellLabs, Sydney, Australia). Filter wheels were air-dried for 4–12 hours before being sealed in ziplock bags with desiccant sachets. These were refrigerated for up to one week and then stored at −20°C before shipping at ambient temperature to LSHTM.

The proportion of children with infection but without active trachoma was low (0.4%) during pre-MDA mapping, therefore swabs were only collected from those with TF during this survey. Swabs were passed three times with a 120°-turn between each pass over the right conjunctiva of children aged 1–9 years who had signs of TF and/or TI in the right eye. Swabs were refrigerated for up to one week and then stored at −20°C before shipping to LSHTM on dry ice for processing. The proportion of active trachoma cases in study villages before MDA was extracted from the full baseline dataset and presented here for comparison.

### Serological and nucleic acid testing

Anti-Ct Pgp3 antibody titre was assessed using enzyme-linked immunosorbent assay (ELISA)^1920^. Optical density (OD) at 450 nm was measured using SpectraMax M3 photometric plate reader (Molecular Devices, Sunnyvale, USA) then normalised to a 20% dilution of presumed-positive standard in presumed-negative standard.

DNA was extracted from swabs with the QIAamp DNA mini kit (Qiagen, Manchester, UK). We tested samples for *Homo sapiens* ribonuclease subunit (RPP30 endogenous control) and open reading frame 2 of the *Ct* plasmid (diagnostic target) using a droplet digital PCR assay^21^ with minor modifications^22^.

### Data analysis

All data analyses were conducted using R 3.2.3^23^. Pre- and post-MDA proportions were compared using Wilcoxon’s rank sum test. ddPCR tests for current ocular *Ct* infection were classified into negative and positive populations according to methods described previously ^21^. ELISAs for antibodies to *Ct* were classified as negative or positive using an expectation-maximisation finite mixture model ^24^. Using this method, the threshold normalised OD value for positivity was 0.7997.

## Results

### Study demographics

In total, 1511 people (46.3% male; 466 1–9 year-olds) aged 1 year and over were examined in 382 households from the 13 selected study villages. By comparison, the pre-MDA survey of the same villages yielded 1534 people (490 1–9 year-olds) in 394 households. Data on non-participation were not collected in June 2015, but the number enrolled was similar to that for the pre-MDA survey, suggesting a similar participation rate (~90%) on both occasions. In this study, there was a mean of 4 people per household aged 1 year and over, and a mean of 1.2 children per household aged 1–9 years, which, after accounting for non-participants, are similar to the means in the 2009 Solomon Islands National Census (4.9 people of any age and 1.4 children aged 1–9 years per household in Temotu, 4.4 people of any age and 1.1 people aged 1–9 years per household in Rennell & Bellona).^25^

### Active trachoma and TT

Prior to MDA, there were 165/489 (33.7%) cases of TF in either eye and 1/489 (0.2%) case of TI in those aged 1–9 years in study villages^13^. Following MDA, we observed 66/466 (14.2%) cases of TF and no cases of TI in either eye, representing a decrease in TF of 58% (p<0.0001). A similar pattern was observed in right eyes considered alone – the eyes from which swabs were collected if indicated (Table 2). 56% of TF cases following MDA were bilateral. No cases of TT were identified during this study.

**Table 2.** Population characteristics of study populations before and after MDA, 13 selected communities of Temotu and Rennell & Bellona Provinces, Solomon Islands.

| Characteristic | Pre-MDA<br>(October–<br>November 2013;<br><sup>13</sup>) | Post-MDA<br>(June–July<br>2015: this<br>study) | p-value* |
| --- | --- | --- | --- |
| Number examined, all ages | 1534 | 1511 | - |
| Number examined aged 1–9 years | 490 | 466 | - |
| Number of households enrolled | 394 | 382 | - |
| % male of those examined | 46.5 | 46.3 | 0.836 |
| TF in either eye in 1–9 year olds | 33.7% | 14.2% | <0.0001 |
| Active trachoma in swabbed eye<br>(right eye field assessment) | 160 (32.7%) | 61 (13.1%) | <0.0001 |
| *Ct* infection in those aged 1–9 years | 5 (3.1%) | 6 (9.8%) | 0.08 |
| Median *Ct* infection load in positive specimens<br>(plasmid copies/swab) | 51,880 | 104,100 | 0.219 |
| *Ct: Chlamydia trachomatis* |  |  |  |

\* t-test

In the two enrolled villages in Rennell & Bellona, a slight increase in the prevalence of TF in either eye in those aged 1–9 years following MDA was noted, but it was not statistically significant (11/60 [17.9%] before MDA to 14/78 [18.3%] after MDA; p=0.956). In contrast, in the 11 enrolled villages of Temotu, a substantial decrease in TF (from 155/430 [36.0%] before MDA to 52/388 [13.4%] after MDA; p<0.0001) was observed.

### Trachomatous scarring

Of the right eye photographs collected, 1440/1511 (95.3%) were suitable for grading. 188/1440 (13.1%) photographs were graded as C>0, of which 127 were C1, 53 were C2 and 8 were C3. Exemplars of scarring that resembled normal trachoma phenotypes are shown in Figures 3A and 3B. The photo-graders noted that some conjunctivae met the criteria for C3, with clear bands of scarring, but also showed clear features not typically associated with trachomatous pathology. In some cases, these were characterised by boundaries demarcating heavily scarred from apparently healthy conjunctiva (Figure 3C and 3D). Photograders noted atypical scars in 4/53 (7.5%) C2 cases and 3/8 (37.5%) C3 cases. Of the scarred eyelids classed as typical for trachoma, 36/54 (67%) were seropositive, whereas 2/7 (29%) of those classed as ‘atypical’ were seropositive, although the difference in proportions was not significant (chi-squared test p=0.123).

**Figure 1:**
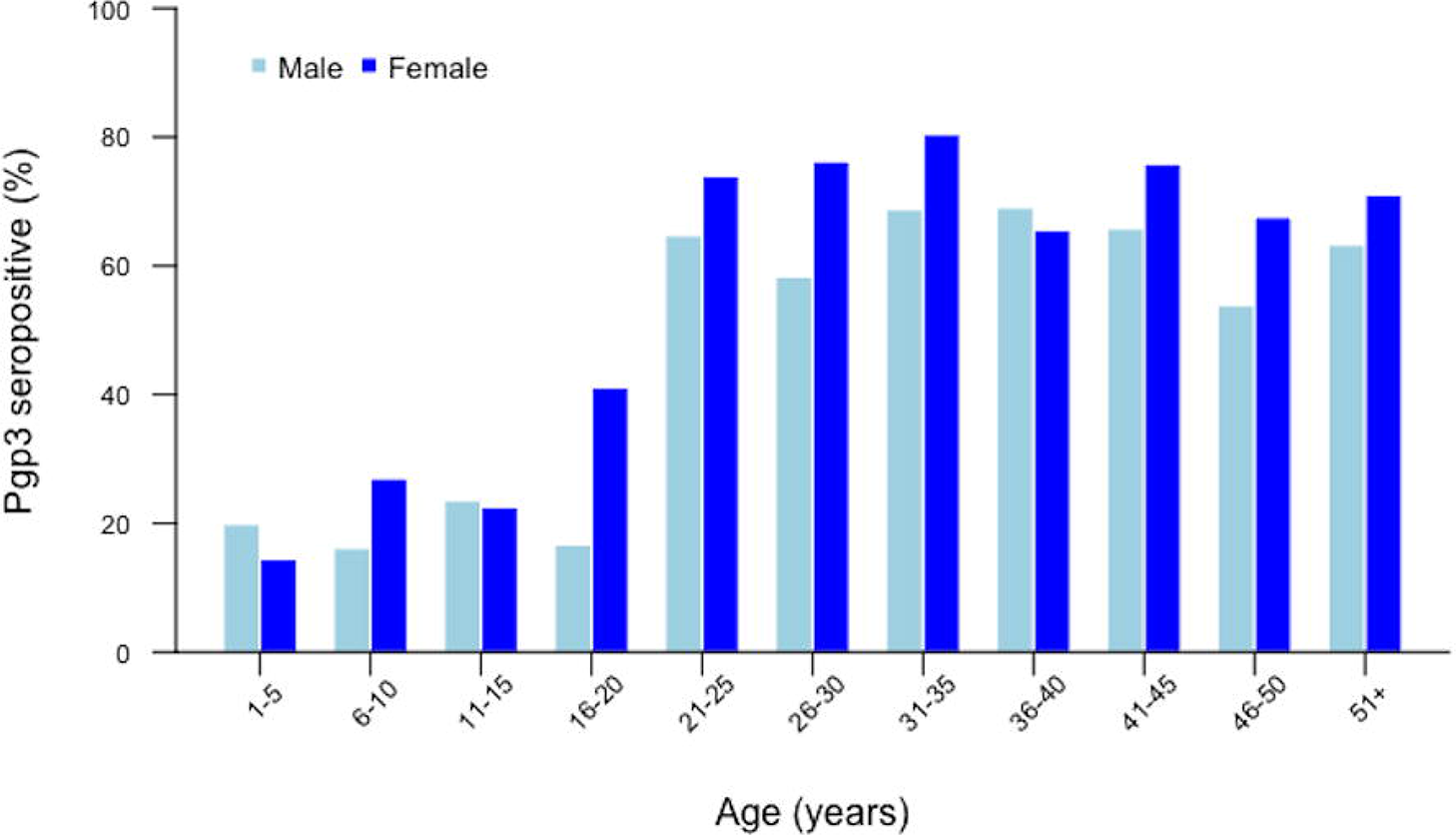
Age-specific seroprevalence of anti-Pgp3 antibodies from dried blood spots collected from 13 selected communities of Temotu and Rennell & Bellona Provinces, Solomon Islands, June-July 2015.

**Figure 2:**
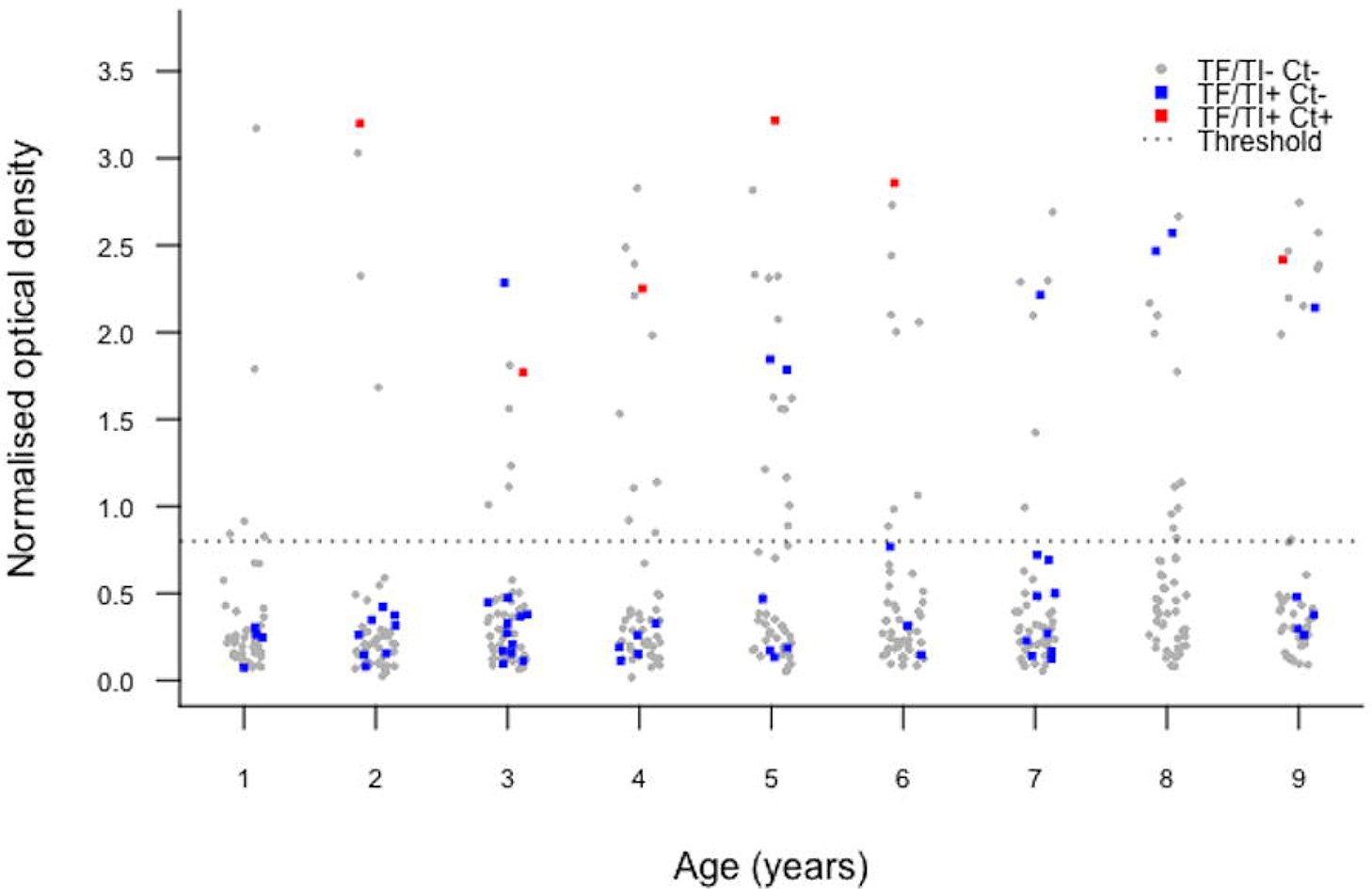
Anti-Pgp3 antibody titre from children aged 1–9 years in 13 selected communities of Temotu and Rennell & Bellona Provinces, Solomon Islands, June-July 2015. Those with TF and/or TI but no infection are highlighted in blue, those with TF and/or TI and infection are red and those with neither TF nor TI nor infection are grey.

**Figure 3:**
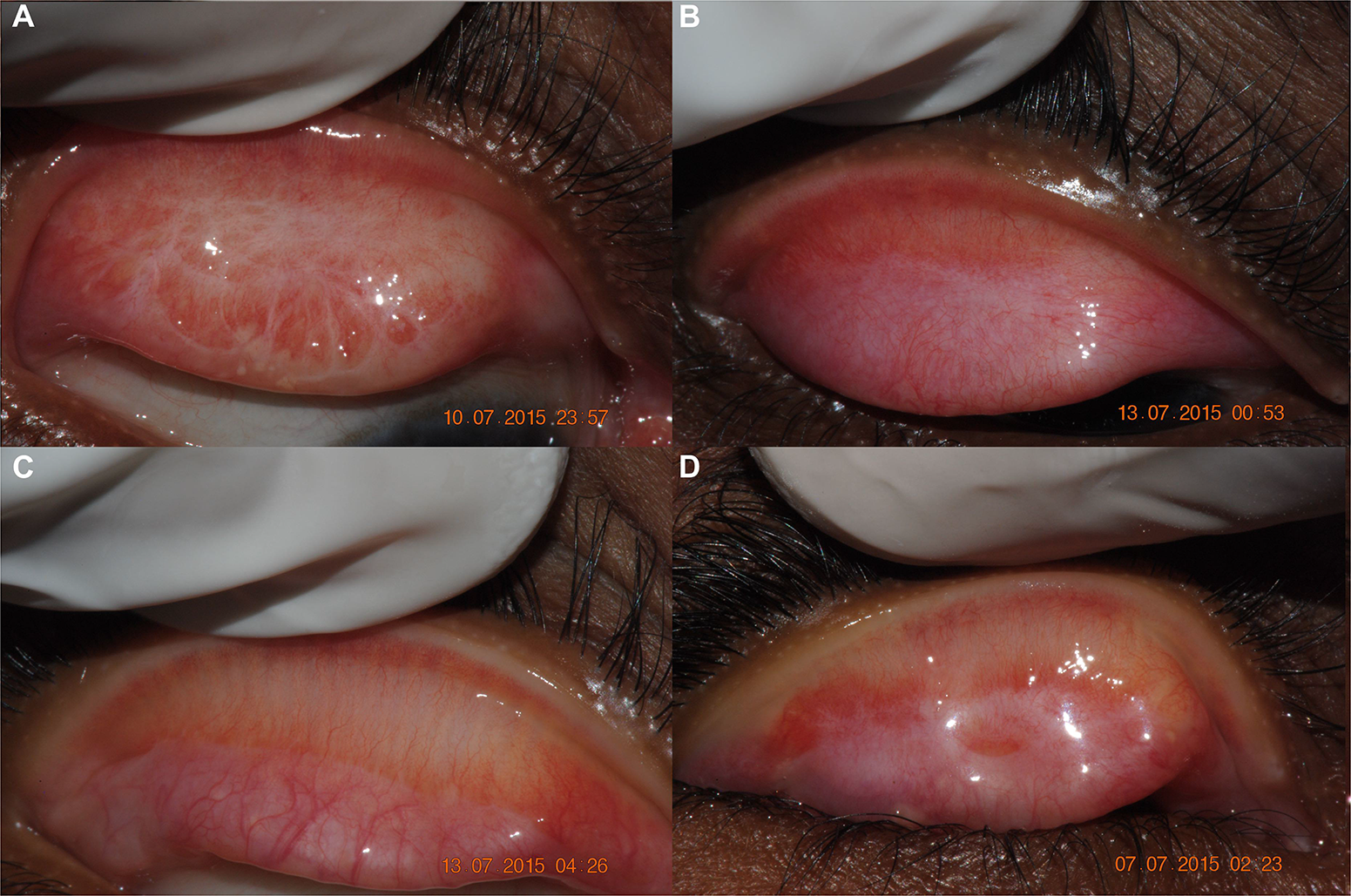
Photographs taken of residents in 13 selected communities of Temotu and Rennell & Bellona Provinces, Solomon Islands, June–July 2015. (**A** and **B**) Conjunctivae with features meeting the criteria for C3 characteristic of trachoma. (**C** and **D**) Conjunctivae with features meeting the criteria for C3 but thought not to be trachomatous in origin. Study IDs were SB113564, SB108878, SB107613 and SB108669.

The age-specific prevalence of scarring is shown in Figure 4. Of 435 photographs graded from children aged 1–9 years, 25 (5.7%) were graded as C>0. In 311 adults aged >40 years who were examined, 74 (23.8%) had C>0 (65 cases of C1, 9 cases of C2, 0 cases of C3). Of 8 cases of C3 in the population, 4 (50%) were in children aged 1–9 years, although 2 of these were classed as ‘atypical’.

**Figure 4:**
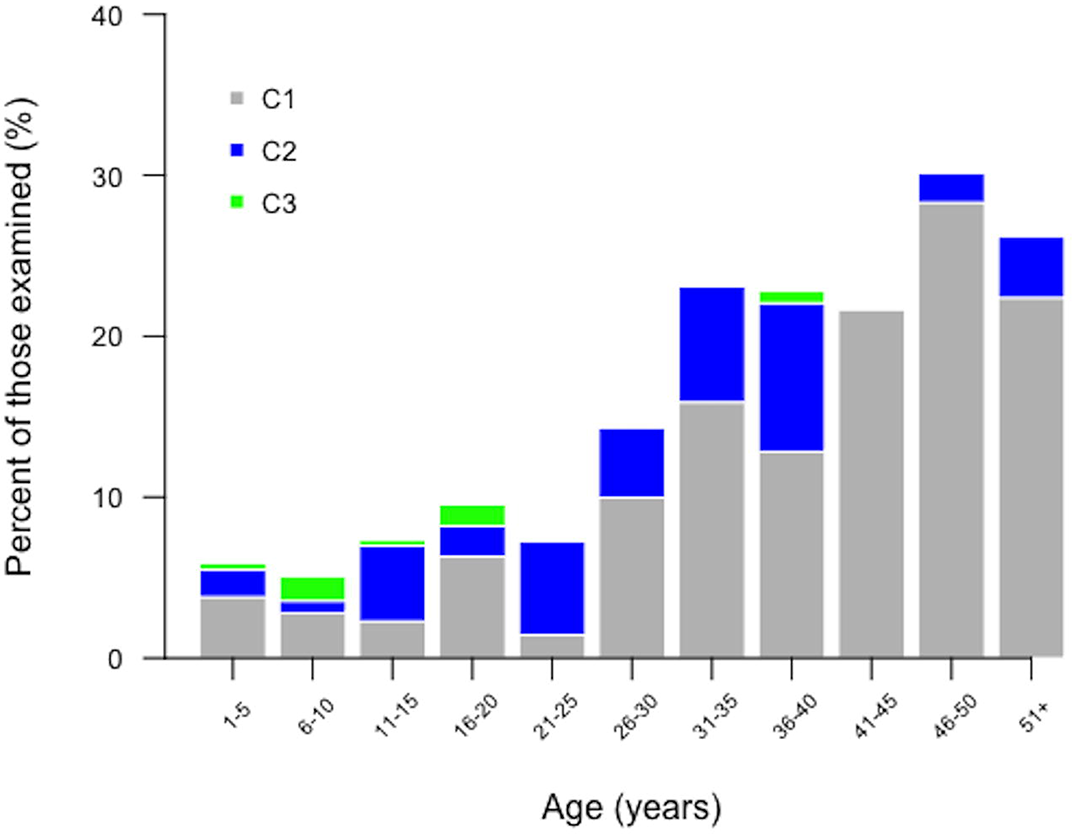
Age-specific prevalence of scarring (defined as C > 0), as identified by photograph grading, in 13 selected communities of Temotu and Rennell & Bellona Provinces, Solomon Islands, June-July 2015.

The proportion of people with C1 increased with age (logistic regression p<0.0001) but the proportion of people with more severe scarring (C2 or C3) did not increase with age (logistic regression p=0.149). There was no significant association between having C>0 and gender (chi-squared test p=0.80). In Rennell & Bellona, 25/225 (11.1%) of photos were graded C>0, whereas in Temotu, 163/1215 (13.4%) of photos were graded C>0; the difference in scarring between provinces was not significant (Wilcoxon Rank Sum test p=0.289).

### Anti-Pgp3 serology

Dried blood spots were collected from 1499/1511 (99.2%) people of all ages during the post-MDA survey; the other 12 people declined finger-prick. Overall, 633/1499 (42.2%) people were considered to be seropositive. In children aged 1–9 years, the prevalence of anti-Pgp3 antibodies was 83/462 (18.0%); in 1-year-olds alone, it was 5/47 (10.6%). The mean seroprevalence in those aged 6–10 years was not significantly higher than in those aged 1–5 years (20.3% compared to 16.6%, Wilcoxon rank sum p=0.276). The largest increase in seroprevalence was observed between those aged 16–20 years and 21–25 years when the seroprevalence rose significantly from 30.4% to 71.6% (Wilcoxon rank sum p<0.0001). Of those aged over 25 years, 67.4% were seropositive (Figure 1). In the 16–20-year-old age group, the prevalence of seropositivity amongst females was higher than in males (13.9% versus 41.1%, Wilcoxon rank sum test p<0.0001). The seroprevalence among children in Rennell & Bellona was significantly higher than that in Temotu (38.5% versus 13.8%; chi-squared p<0.0001).

There was no association between seropositivity and signs of TF in children aged 1–9 years (19.7% seropositive in those with TF in either eye compared to 17.7% seropositive in those without TF, p=0.581). In those younger than the median self-reported age of sexual debut (18 years ^26^), there was no association between C grade and anti-Pgp3 OD (linear regression adjusted for age and gender p=0.453) or anti-Pgp3 positivity (logistic regression adjusted for age and gender p=0.472).

### Ocular *Chlamydia trachomatis* infection

Swabs from all 61 children aged 1–9 years with TF in the right eye that were tested for *Ct* had a positive endogenous control result; the median load of the human RPP30 target was 83,000 copies, equivalent to over 40,000 conjunctival cells. In this study, 6/61 (9.8%) of children with active trachoma had *Ct* infection. Of the 6 specimens from children positive for *Ct,* the median load was 104,100 plasmid copies/swab. We previously showed that before MDA, 5/160 (3.1%) of those with active trachoma in study villages had evidence of infection with *Ct.* The median pre-MDA load of *Ct* infections in these villages was 51,880 plasmid copies/swab.^13^ Neither the difference between the pre- and post-MDA *Ct* prevalence nor pre- and post-MDA *Ct* load was statistically significant (Wilcoxon rank sum test p=0.08 and p=0.22, respectively). The relationship between *Ct* infection, signs of trachoma and seropositivity was examined in children aged 1–9 years and is summarised in Table 3. All six cases of active trachoma in which infection was also detected were in seropositive individuals (Figure 2). All study villages had at least one case of TF, but infections were limited to three of the 13 villages studied. Two villages in Rennell & Bellona housed 5/6 *Ct* infections identified during this study.

**Table 3.** Serological status compared to other tests for trachoma, 13 selected communities of Temotu and Rennell & Bellona Provinces, Solomon Islands, June-July 2015

| Comparator |  | 1–9 year-olds |  |  | ≥10 year-olds |  |  |
| --- | --- | --- | --- | --- | --- | --- | --- |
|  |  | Seronegative | Seropositive | Total | Seronegative | Seropositive | Total |
| Ct infection by ddPCR* | Positive | 0 | 6 | 6 | - | - | - |
|  | Negative | 48 | 7 | 55 | - | - | - |
| TF | Positive | 53 | 13 | 66 | 13 | 9 | 22 |
|  | Negative | 326 | 70 | 396 | 474 | 541 | 1015 |
| TI | Positive | 0 | 0 | 0 | 0 | 1 | 1 |
|  | Negative | 379 | 83 | 462 | 487 | 549 | 1036 |
| Scarring | C0 | 333 | 77 | 410 | 414 | 418 | 832 |
|  | C1 | 15 | 1 | 16 | 36 | 75 | 111 |
|  | C2 | 3 | 2 | 5 | 16 | 32 | 48 |
|  | C3 | 3 | 1 | 4 | 1 | 3 | 4 |

Ct: *Chlamydia trachomatis*; ddPCR: droplet digital polymerase chain reaction; TF: trachomatous inflammation—follicular;
TI: trachomatous inflammation—intense.
\* presence or absence of infection only assessed in children with TF and/or TI.

## Discussion

The Solomon Islands, along with other Pacific Island states, has been identified as trachoma-endemic based on moderately high province-level prevalences of TF. Whilst measures for trachoma elimination have already been deployed in Temotu and Rennell & Bellona, we have previously noted that TI, ocular *Ct* infection and late-stage disease (TT) are rare^13^. If the findings from these study villages were replicated throughout the district, TF would still be sufficiently prevalent to warrant intervention. However, using a suite of non-TF tools (clinical photography for evaluation of conjunctival scarring, nucleic acid infection testing and serological testing), we have demonstrated here that ocular *Ct* is scarce and is not being widely transmitted, and that TF is not concurrent with prevalent severe scarring or TT in this population.

We would not expect to see large numbers of individuals with TF who have not previously been infected with *Ct*, but in this population, the majority (80.3%) of individuals with TF were seronegative, and participants with TF were no more likely to be seroreactive to Pgp3 than their peers without TF. We found a small and non-significant increase in age-specific seroprevalence between young children (0–5 years) and older children (6–10 years), which suggests that there is limited horizontal transmission of *Ct* strains among children. This is concordant with our previous data, which suggested that although ocular *Ct* strains are present in the Solomon Islands, they are rare^13^. The increase in seropositivity with age in this group was modest compared with that seen in hyperendemic villages of the United Republic of Tanzania, where seropositivity has been observed to increase from approximately 25% to 94% between the ages of 1 and 6 years.^19^ In the current dataset, there was a rapid increase in age-specific seroprevalence around the age of 18 years, the self-reported median age of sexual debut in a nearby population^27^. The prevalence of urogenital *Ct* infection is known to be high in women attending antenatal clinics in the Solomon Islands ^27^, which may account for the high seroprevalence in adults, and exposure during parturition may also be a major contributor to the 10% of 1-year-olds in our study who had evidence of prior Pgp3 exposure.^28^

Antibodies to Pgp3 have recently been suggested for monitoring *Ct* transmission in trachoma programmes^10,29^, but there is still much to learn about the dynamics of these responses. For example, it is not clear whether anti-Pgp3 responses are detectable in all individuals previously infected with *Ct,* or if multiple exposures are required to develop sustained anti-Ct responses. In this study, only 19.7% of children with TF were positive for anti-Pgp3 antibodies, but all six children with current infection were seropositive. There was also a high prevalence of Pgp3 reactivity in adults living in Temotu and Rennell & Bellona, a proportion of whom are likely to have had a previous urogenital *Ct* infection^27^. While seroreversion due to clearance of infection by MDA is a possible explanation for the low seroprevalence and absence of association of anti-Pgp3 antibodies with TF, there is currently no evidence for complete seroreversion for Pgp3-specific antibodies^9,19^ after clearance of infection. We are therefore confident that Pgp3 is an appropriate antigen for serosurveillance in this population.

Analysis of the age-specific prevalence of conjunctival scarring illustrated that, while the proportion of people with mild scars increased with age, the proportion of those with more extensive or eyelid-distorting scars did not increase with age. Contrary to what might be expected in a trachoma-endemic community^30^, no eyelid-distorting scars were found in 311 adults >40 years. There does appear to be severe scarring among children, but some cases are atypical (Figure 3) and are in children who lack Pgp3 reactivity (Table 3), so other causes of scarring may be contributing to this (presumed) ongoing incidence. There are a number of inflammatory conditions (e.g. adenoviral, acute haemorrhagic or membranous conjunctivitis) which may result in conjunctival scarring, although the pathology, incidence and prevalence of these are poorly understood.^31^ It is currently unclear whether the TF that we observed is directly linked to conjunctival scarring in this setting. Currently in Temotu and Rennell & Bellona, the low prevalence of severe scars suggests that the proportion of the population at risk of developing TT is very low, although we cannot determine how this might change temporally.

Prior to intervention with MDA, more than 15% of children living in the selected villages had TF. The 13 communities included here were the most highly endemic of those surveyed in Temotu and Rennell & Bellona during the GTMP. In this study, we showed that the burden of TF in many of these villages dropped significantly following a single round of MDA, but still remains above the threshold for continued intervention. The drop in clinical disease was not reflected by a drop in ocular *Ct* in children with TF, which increased, albeit statistically insignificantly. From interventions in other settings, 6 months after a single round of MDA we might expect TF prevalence to drop by approximately 50%, given 80% population coverage^32,33^. Azithromycin has anti-inflammatory and broad-spectrum antibiotic effects, which may help explain the observed decrease in clinical disease. We observed regional variation across the study villages. Compared to Temotu, we noted more children were seropositive, more children with TF had infection, and MDA did not have as significant an impact on TF levels in Rennell & Bellona. Our survey was not prospectively designed to assess these differences, and the subgroup size in Rennell & Bellona precludes more detailed analysis. Temotu is much more similar to the rest of the Solomon Islands in terms of the geology of the islands, and the lifestyle and ethnicity of the majority of the inhabitants. Further studies on the localisation of trachoma in the islands are warranted.

There were limitations to our study. Firstly, due to the remoteness of the islands studied, both blood and swab samples were stored for up to 24 hours at room temperature (commonly around 25–30°C in the Solomon Islands) and up to one week at 4°C prior to freezing. DNA and antibodies may degrade if stored for prolonged periods at high temperatures, however, several studies have now shown that short- and long-term storage of swabs at room temperature will not cause significant loss of diagnostic signal in any but the lowest load (and therefore least relevant to transmission) samples^22,34–36^. Antibodies too are very stable even after short-term refridgeration^37^. Secondly, some commentators have criticised the sensitivity of the ddPCR diagnostic used in this study^38^. The technique was evaluated against Amplicor and demonstrated only to fail diagnostically in the lowest load samples^21^. Similar protocols were used in other studies in the Pacific and elsewhere in the world and were able to detect infections at high prevalence^39–41^. Finally, only one antigen was tested for during serological testing using an assay which is still under evaluation. Although Pgp3 is emerging as the marker of choice for *Ct* serosurveillance^10,42^, verification of these data by testing for an independent immunodominant antigen such as major outer membrane protein with microimmunofluorescence may increase our confidence in the findings.

The complex, multistage nature of trachoma makes it difficult to predict the outcome of a given intervention^43^. Data from cross-sectional surveillance tools used in isolation can be hard to interpret, especially given the prolonged persistence of TF after clearance of infection^44^. Some features of conjunctivitis in the Solomon Islands resemble trachoma, particularly the prevalent follicular inflammation and some of the severe conjunctival scarring. Crucially, these clinical features were not co-endemic with TT at a prevalence that indicates a public health problem. In this setting, tests for infection gave a better indication of the public health threat from trachoma than TF. A combined approach in which various age-specific markers of trachoma are assessed together across the complete age range of the population, may prove useful for prioritising areas for intervention where the prevalence of TF alone does not coherently reflect trachoma’s public health importance.

Contrary to the WHO recommendation for treatment based solely on prevalence of TF, our data provide compelling evidence that trachoma is not a public health problem in these villages. Whilst there have been substantial collateral benefits to local residents from having received MDA (such as on genital *Ct^45^* and yaws^46^), further rounds of azithromycin MDA do not appear to be indicated for the purposes of trachoma elimination. As the positive predictive value of TF decreases globally, other countries may emerge where TF is not reflective of threat to vision. WHO recommendations for implementation of MDA and the SAFE strategy should be reviewed in the light of this evidence.

## Acknowledgements

We are grateful to the survey participants in Temotu and Rennell & Bellona, to Andrew Velaio for logistical support in the field, and to Leslie Sui, Charles Russell and Suzanne Tetepitu for helping to conduct the field work. The findings and conclusions in this report are those of the authors and do not necessarily represent the official position of the Centers for Disease Control and Prevention.

## Financial statement

The field and laboratory costs were funded by the Fred Hollows Foundation (1041). RMRB and AWS were funded by a Wellcome Trust Intermediate Clinical Fellowship to AWS (098521).

OS and KJ were employed by the Solomon Islands Ministry of Health and Medical Services for the duration of the survey.

DLM receives funding through the US Agency for International Development through an interagency agreement with CDC.

MJB was funded by the Wellcome Trust (098481/Z/12/Z).

ChR is supported by the Wellcome Trust Institutional Strategic Support Fund (105609/Z/14/Z).

The funders had no role in the design, execution or publication of this study.

## Author Contributions

Conceived and designed the study: RMRB, OS, DLM, AWS, DCWM, ChR.

Performed the fieldwork: RMRB, OS, KJ Provided training and reagents: DLM Performed the experiments: RMRB, MJB, ChR Analysed the data: RMRB, MJB, ChR

Wrote the manuscript: RMRB, ChR

Revised and approved the manuscript: RMRB, OS, KJ, DLM, MJB, AWS, DCWM, ChR

## References

1. Bourne, R. A. et al. Causes of vision loss worldwide, 1990–2010: a systematic analysis. Lancet Glob. Heal. 1, e339–49 (2013).

2. Wolle, M. A., Muñoz, B. E., Mkocha, H. & West, S. K. Constant ocular infection with Chlamydia trachomatis predicts risk of scarring in children in Tanzania. Ophthalmology 116, 243–7 (2009).

3. West, S. K., Muñoz, B., Mkocha, H., Hsieh, Y. H. & Lynch, M. C. Progression of active trachoma to scarring in a cohort of Tanzanian children. Ophthalmic Epidemiol. 8, 137–44 (2001).

4. King, J. et al. Impact of the SAFE strategy on trachomatous scarring among children in Ethiopia. in Abstracts of the 9th European Congress on Tropical Medicine and International Health. 6–10 September (ed. Tropical Medicine and International Health) 20, 240 (2015).

5. Hu, V. H., Holland, M. J. & Burton, M. J. Trachoma: protective and pathogenic ocular immune responses to Chlamydia trachomatis. PLoS Negl. Trop. Dis. 7, e2020 (2013).

6. Kari, L. et al. Antibody signature of spontaneous clearance of Chlamydia trachomatis ocular infection and partial resistance against re-challenge in a nonhuman primate trachoma model. PLoS Negl. Trop. Dis. 7, e2248 (2013).

7. Ghaem-Maghami, S. et al. Mucosal and systemic immune responses to plasmid protein pgp3 in patients with genital and ocular Chlamydia trachomatis infection. Clin. Exp. Immunol. 132, 436–442 (2003).

8. Comanducci, M. et al. Humoral immune response to plasmid protein pgp3 in patients with Chlamydia trachomatis infection. Infect. Immun. 62, 5491–5497 (1994).

9. Horner, P. et al. C. trachomatis Pgp3 antibody prevalence in young women in England, 1993–2010. PLoS One 8, e72001 (2013).

10. Goodhew, E. B. et al. CT694 and pgp3 as serological tools for monitoring trachoma programs. PLoS Negl. Trop. Dis. 6, e1873 (2012).

11. World Health Organization. Report of the 3rd Global Scientific Meeting on Trachoma. 19–20 July. (2010).

12. Ramadhani, A. M., Derrick, T., Macleod, D., Holland, M. J. & Burton, M. J. The Relationship between Active Trachoma and Ocular Chlamydia trachomatis Infection before and after Mass Antibiotic Treatment. PLoS Negl. Trop. Dis. 10, e0005080 (2016).

13. Butcher, R. M. R. et al. Low Prevalence of Conjunctival Infection with Chlamydia trachomatis in a Treatment-Naïve Trachoma-Endemic Region of the Solomon Islands. PLoS Negl. Trop. Dis. 10, e0004863 (2016).

14. Martin, D. L. et al. Serological Measures of Trachoma Transmission Intensity. Sci. Rep. 5, 18532 (2015).

15. Kirkwood, B. & Sterne, J. A. in Essentail Medical Statistics 413–428 (Blackwell Publishing Ltd, 2003).

16. Thylefors, B., Dawson, C. R., Jones, B. R., West, S. K. & Taylor, H. R. A simple system for the assessment of trachoma and its complications. Bull. World Health Organ. 65, 477–83 (1987).

17. Solomon, A. W. et al. The Global Trachoma Mapping Project: Methodology of a 34-Country Population-Based Study. Ophthalmic Epidemiol. 22, 214–25 (2015).

18. Dawson, C. R., Jones, B. R., Tarizzo, M. L. & World Health Organization. Guide to trachoma control in programmes for the prevention of blindness. Geneva, Switzerland. (1981).

19. Martin, D. L. et al. Serology for Trachoma Surveillance after Cessation of Mass Drug Administration. PLoS Negl. Trop. Dis. 9, e0003555 (2015).

20. Cocks, N. et al. Community seroprevalence survey for yaws and trachoma in the Western Division of Fiji. Trans. R. Soc. Trop. Med. Hyg. 110, 582–587 (2016).

21. Roberts, C. H. et al. Development and Evaluation of a Next-Generation Digital PCR Diagnostic Assay for Ocular Chlamydia trachomatis Infections. J. Clin. Microbiol. 51, 2195–203 (2013).

22. Macleod, C. K. et al. Low prevalence of ocular Chlamydia trachomatis infection and active trachoma in the Western Division of Fiji. PLoS Negl. Trop. Dis. 10, e0004798 (2016).

23. R Core Team. R: A Language and Environment for Statistical Computing. R Foundation for Statistical Computing (2014). Available at: http://www.r-project.org.

24. Migchelsen, S. J. et al. Defining Seropositivity Thresholds for Use in Trachoma Elimination Studies. PLoS Negl. Trop. Dis. 11, e0005230 (2017).

25. Solomon Island Government. Report on 2009 population and housing census. (2011).

26. Marks, M. et al. Prevalence of sexually transmitted infections in female clinic attendees in Honiara, Solomon Islands. BMJ Open In Press, (2015).

27. Marks, M. et al. Prevalence of sexually transmitted infections in female clinic attendees in Honiara, Solomon Islands. BMJ Open 5, (2015).

28. Schachter, J. et al. Propsective study of chlamydial infection in neonates. Lancet 2, 377–380 (1979).

29. Goodhew, E. B. et al. Longitudinal analysis of antibody responses to trachoma antigens before and after mass drug administration. BMC Infect. Dis. 14, 216 (2014).

30. Wolle, M. A., Muñoz, B., Mkocha, H. & West, S. K. Age, sex, and cohort effects in a longitudinal study of trachomatous scarring. Invest. Ophthalmol. Vis. Sci. 50, 592–6 (2009).

31. Faraj, H. G. & Hoang-Xuan, T. Chronic cicatrizing conjunctivitis. Curr. Opin. Ophthalmol. 12, 250–7 (2001).

32. Yohannan, J. et al. Can we stop mass drug administration prior to 3 annual rounds in communities with low prevalence of trachoma?: PRET Ziada trial results. JAMA Ophthalmol. 131, 431–6 (2013).

33. Burton, M. J. et al. Profound and sustained reduction in Chlamydia trachomatis in The Gambia: a five-year longitudinal study of trachoma endemic communities. PLoS Negl. Trop. Dis. 4, 10 (2010).

34. van Dommelen, L. et al. Influence of temperature, medium, and storage duration on Chlamydia trachomatis DNA detection by PCR. J. Clin. Microbiol. 51, 990–2 (2013).

35. Gaydos, C. A. et al. Can mailed swab samples be dry-shipped for the detection of Chlamydia trachomatis, Neisseria gonorrhoeae, and Trichomonas vaginalis by nucleic acid amplification tests□? Diagn. Microbiol. Infect. Dis. 73, 16–20 (2012).

36. Dize, L., Gaydos, C. A., Quinn, T. C. & West, S. K. Stability of Chlamydia trachomatis on storage of dry swabs for accurate detection by nucleic acid amplification tests. J. Clin. Microbiol. 53, 1046–1047 (2014).

37. Corran, P. H. et al. Dried blood spots as a source of anti-malarial antibodies for epidemiological studies. Malar. J. 7, 195 (2008).

38. Schachter, J. Will droplet digital PCR become the test of choice for detecting and quantifying ocular Chlamydia trachomatis infection? Maybe not. Expert Rev. Mol. Diagn. 13, 789–92 (2013).

39. Mueller, A. J. Assessment of the Prevalence of Trachoma on Kiritimati, Republic of Kiribati. (2016).

40. Derrick, T. et al. Can corneal pannus with trachomatous inflammation - follicular be used in combination as an improved specific clinical sign for current ocular Chlamydia trachomatis infection? Parasit. Vectors 9, (2016).

41. Butcher, R. et al. Reduced-cost Chlamydia trachomatis-specific multiplex real-time PCR diagnostic assay evaluated for ocular swabs and use by trachoma research programmes. J. Microbiol. Methods S0167–7012, 30097–0 (2017).

42. Wills, G. S. et al. Pgp3 antibody enzyme-linked immunosorbent assay, a sensitive and specific assay for seroepidemiological analysis of Chlamydia trachomatis infection. Clin. Vaccine Immunol. 16, 835–43 (2009).

43. Liu, F. et al. Short-term Forecasting of the Prevalence of Trachoma: Expert Opinion, Statistical Regression, versus Transmission Models. PLoS Negl. Trop. Dis. 9, (2015).

44. West, S. S. K. et al. Can We Use Antibodies to Chlamydia trachomatis as a Surveillance Tool for National Trachoma Control Programs? Results from a District Survey. PLoS Negl. Trop. Dis. 10, e0004352 (2016).

45. Marks, M. et al. Mass drug administration of azithromycin for trachoma reduces the prevalence of genital Chlamydia trachomatis infection in the Solomon Islands. Sex. Transm. Infect. (2016). doi:10.1136/sextrans-2015-052439

46. Marks, M. et al. Impact of community mass treatment with azithromycin for trachoma elimination on the prevalence of yaws. PLoS Negl. Trop. Dis. 9, e0003988 (2015).

